# Development of non-destructive methods to estimate functional traits and field evaluation in tea plantations using a smartphone

**DOI:** 10.1101/2020.03.17.994897

**Authors:** Esack Edwin Raj, Rajagobal Raj Kumar, Ashokraj Shanmugam, Balakrishnan Radhakrishnan

**Affiliations:** Plant Physiology and Biotechnology Division, UPASI Tea Research Foundation, Tea Research Institute, Nirar Dam PO, Valparai 642 127, Tamil Nadu, INDIA

**Keywords:** Chlorophyll, Disease assessment, Image segmentation, Nitrogen, SPAD meter

## Abstract

Smartphones are equipped with various types of sensors which make them a promising tool to assist diverse digital farming tasks because of their mobility, cost, accessibility, and computing power allow us to perform real-time practical applications. This paper presents the utilization of various non-destructive methods of nutrient and disease classification techniques using smartphone collected images, processed through various image segmentation algorithms. Both *in vivo* and *in vitro* estimations shows comparable results with both chlorophyll and nitrogen contents of a crop shoot. Moreover, the correlation between SPAD measured values and nitrogen of crop shoot showed a significant linear association (R^2^ =0.7309), revealing the potency of *in vivo* observation for prediction of actual chlorophyll content in tea crop. SPAD values and yield have a strong linear relationship (R^2^ =0.7103), in which SPAD-meter performed better detection at very low values. The study concluded that the proposed techniques could be used for automatic detection as well as classification of foliar diseases and nutrients in tea.

## Introduction

In the technology era, industries are shifting from manual to automated solutions for various problems. The developed technology has not only augmented the efficiency, but they also have shortened the time, cost, and labour required to get assured excellence. In agriculture, tea is one of the major economic plantation crops casing a large amount of area and labour. Presently, the industry as a whole facing lot of problems in which nutrient and disease management are primary concerns besides labour shortage, in turn, leads to economic volatility. The idea behind this research is to develop precision non-destructive methods which can work out for the problem of nutrient and disease assessment in the tea crop by examining the image. In the modern world, image processing has fetched it in reality which can be used in any ecological conditions and lead to very momentous, reliable and precise elucidations in problem-solving. Such a study can give a clear idea about nutrition status and its kind, the area of pest incidences, pesticide requirement or just to do what is necessary, etc., for a crop by comparing the colour of a plant’s leaves with the endorsements given by agricultural organizations. The tea industry is highly dependent on productivity where potentially destructive blight diseases can result in ∼35% crop loss. Therefore, the use of disease detection technique plays an important role that could enable planters to take preventive/quarantine measures at the initial stage of disease transition.

Chlorophylls play an important role in the plant as primary photosynthetic pigment composed together with carotenoids exhibits a unique colour appearance which is specific to genotype and even used as a parameter of physiological maturity, nutrient status and quality (Pagola et al. 2009). The chlorophyll content is strongly related to the colour and flavour of the tea is principally determined by crop shoot chemical constituents (Wang et al., 2004) and is positively correlated with the scores for appearance and infused leaf (Wang et al., 2014). Leaf chlorophyll is mostly used as an index to diagnose diseases and nitrogen (N) status in the plants besides minor nutrient symptoms. N is the most required mineral nutrient in tea where the optimum and timely application is crucial in achieving a higher yield. However, an excess application of N in the agricultural environment can lead to water pollution and upon intake, it is transformed into nitrosamines molecules acting as carcinogens (Turner and Rabalais, 1991). Hence, site-specific timely application of N at required quantity with a spatial concern could be a sustainable method to overcome the limitations of traditional replacement theory; however, methods have to be developed for real-time estimation along with cost-effectiveness.

Although disease symptom and nutrient status of the plant can be assessed by human observation, yet initial stage identification, cause for a larger area and their spatial relationship with an associated environmental variable are quite impossible for a common agrarian but very important for disease dynamics. So far, various conventional invasive methods have been practised and modern non-destructive methods are being developed to assess the leaf chlorophyll, nitrogen and disease of various crop plants. Conventional methods of determining the chlorophyll and nitrogen content are destructive chromatographic methods, laborious and are not adaptable to real-time estimation (Gilmore and Yamamoto, 1991). As an alternative, the non-destructive image processing technique is an emerging field of agriculture, enabling real-time measurement using digital images to get the intrinsic characteristics of leaf i.e. colour and texture and are related to geographical information. This method is a very easy, portable, less time consuming, cost-effective and provides accurate information and understanding at a spatial scale. The proposed image processing method in the study deals with trichromatic colours, i.e. RGB and compared to the destructive methods for nitrogen and chlorophyll contents besides disease detection by using various statistical analyses.

## Material and methods

The overview of the methodology proposed in the study depicted in Fig.-1. Initially, the chlorophyll content of UPASI clones was chemically estimated at different seasons for two agricultural years using acetone after recording SPAD-502 chlorophyll meter (Konica Minolta Sensing Inc., Tokyo, Japan). The same leaves were photographed using a smartphone for the estimation of chlorophyll using image processing algorithms. The settings of the camera, includes enabling GPS, sensor light sensitivity (ISO) and exposure time were set in auto mode and the camera could select them based on light conditions. The histogram of leaf image was obtained using ImageJ and extracted RGB values correlated with the SPAD and chemically estimated chlorophyll. Neural network analysis with genetic algorithm was used to estimate the plant pigments by training (70% of the data) and validation (30% of the data). The accuracy of the developed model is based on R^2^, SE and RMSE.

For the development of a non-destructive method of N estimation, a randomised completely block designed (RCBD) experiment was conducted with UPASI-9 clone after two years of pruning. Each treatment replicated three times and each block consisted of ∼100 plants. Experimental plots treated with various levels of Nitrogen (0, 150, 300, 600, 900 and 1600 kg/ha/y in four splits) and images were taken before and after every harvest. Mother leaf collected from respective plots subjected to shade drying and grinding, the N content of the samples was determined by Kjeldahl digestion technique.

**Fig.-1:**
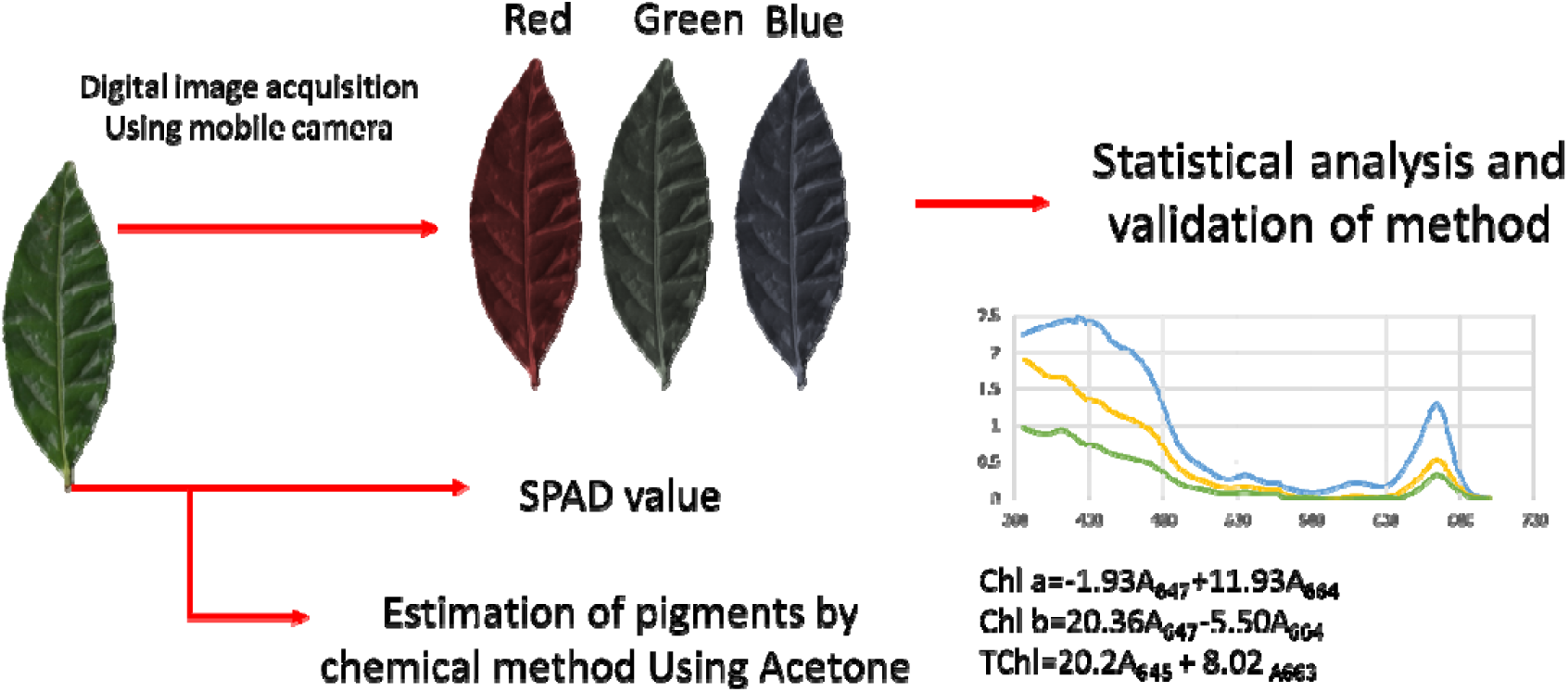
Schematic overview of the proposed study.

Image segmentation algorithm used to classify the diseased with non-diseased parts in the leaf using neural networks with a genetic algorithm. In the first step of fitness computation, the dataset of the pixel is clustered according to nearest respective cluster centres such that each pixel *X*_*i*_ of the colour image is put into the respective cluster with the cluster centre *z*_*j*_ for *j* = 1, 2,…, *K* by the following equations

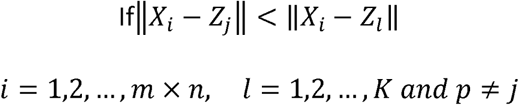

The percentage of disease was spatially interpolated geo-statistically using the kriging method in which the weights are assessed by the autocorrelation to obtain unbiased error and minimum estimation variance. Kriging interpolation uses the attribute values (i.e. nugget, sill, and range values) obtained from semivariogram analysis and was evaluated through a leave-one-out cross-validation approach (Gholizadeh et al., 2012). Maps were created in ArcGIS software.

## Results and Discussion

### Estimation of chlorophyll

Initially, acetone extracted leaf pigments were estimated and the results are correlated with the RGB features extracted from smartphone capture images using ImageJ software was shown in Fig.-2. The correlation between the SPAD estimated pigment values and image processing values shown in Fig.-3. A significant and positive correlation was observed between SPAD meter reading and image algorithm reading at each stage of measurement. The coefficients of determination (R^2^) indicated that there was a minor variation in chlorophyll content which was explained by image processing algorithm reading at each measurement stage of development (Fig.-3b) that ranging from 0.8 to 2% (Table-1). The relationship between image feature and SPAD values indicated that red (R^2^ = 0.830) and green (R^2^ = 0.857) colours are highly related with the SPAD values except for blue colour (R^2^ = 0.006). The result also indicated the SPAD values are negatively correlated with the red (y = −3.650× + 302.054) and green (y = −2.435× + 278.679) while blue is positively related (y = 0.107× + 13.194; Fig.-3a). The results show that the image processing algorithm readings can be used as an effective indicator for evaluating the chlorophyll content of the leaf at a different stage of the crop. Based on this exercise, the reader must conclude that surface chlorophyll measurements are worthwhile by making use of an image processing algorithm.

**Table-1:** Validation of chemically estimated pigments with the SPAD and image extracted values.

| SPAD | R<sup>2</sup> | SE | RMSE | Image | R<sup>2</sup> | SE | RMSE |
| --- | --- | --- | --- | --- | --- | --- | --- |
| Chl-a | 0.862 | 0.322 | 0.568 | Chl-a | 0.951 | 0.061 | 0.247 |
| Chl-b | 0.761 | 0.233 | 0.482 | Chl-b | 0.968 | 0.027 | 0.165 |
| Caro | 0.997 | 0.014 | 0.120 | Caro | 0.931 | 0.024 | 0.156 |
| T-Chl | 0.946 | 0.280 | 0.529 | TChl | 0.938 | 0.101 | 0.317 |
| Xantho | 0.666 | 0.095 | 0.308 | Xantho | 0.974 | 0.007 | 0.086 |

Chl-a: Chlorophyll-a; Chl-b: Chlorophyll-b; Caro: Carotenoid; T-Chl; Total Chlorophyll; Xantho: Xanthophyll; the unit of all the plant pigments are expressed as mg/g; R<sup>2</sup>: Correlation coefficient values; SE: standard error; and RMSE: Root mean sum square error.

**Fig.-2:**
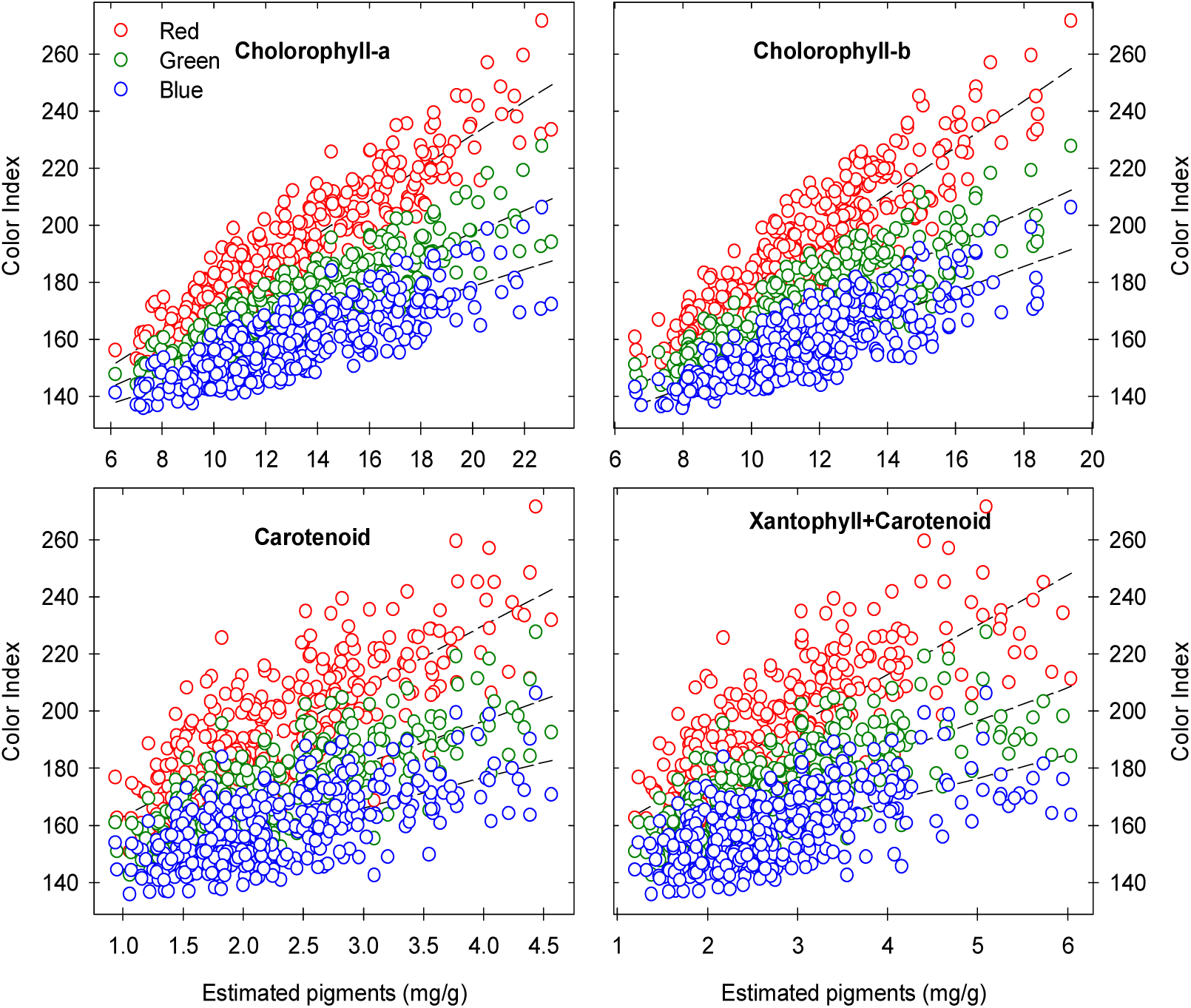
The relationship between RGB features and chemically estimated plant pigments.

**Fig.-3:**
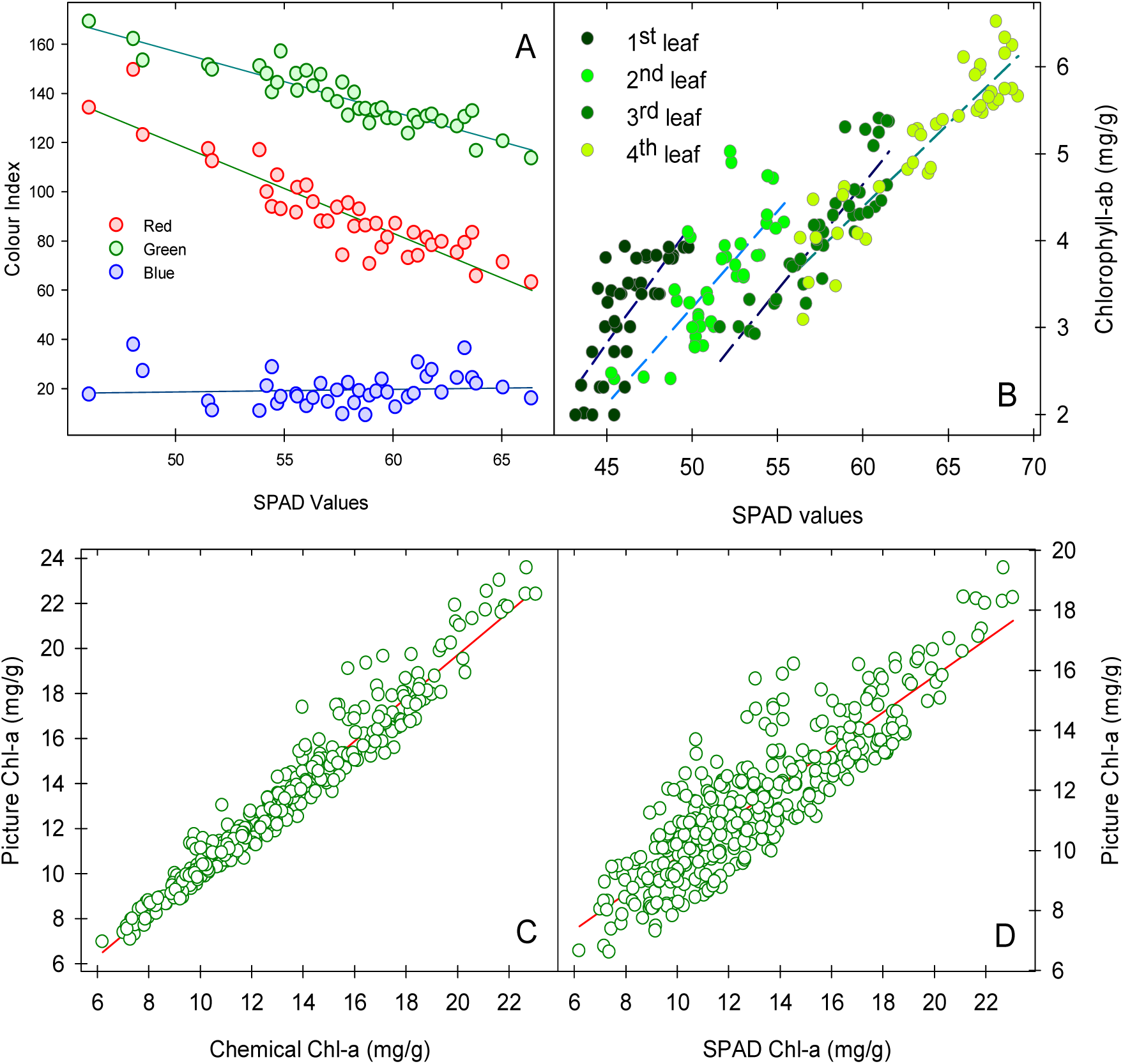
Relationship between SPAD values and chemically estimated values at various stages (A & B). Validation of estimated plant pigments between chemically estimated and image extracted chlorophyll-a (C), similarly with SPAD estimated chlorophyll-a (D).

### Measuring the nitrogen status and its relationship with yield

A regression analysis was made between SPAD readings and leaf N content besides the yield. There was a significant positive linear relationship between both the attributes as displayed in Fig.-4. Regression analysis of the SPAD readings and yield demonstrated that these attributes produced a significant linear relationship R^2^ = 0.7309; y = 85.213× − 3980.2 (Fig.-4a). Similarly, the regression analysis of the SPAD readings and N demonstrated a significant linear relationship R^2^ = 0.7103; y = 0.1868× − 7.8051 (Fig.-4b). Similar results were arrived by Yang et al. (2014), Ramesh et al. (2002) and Swain & Sandip (2010) who established the linear relationships between leaf N content and SPAD readings and highly significant correlations of SPAD readings with the yield in various crops.

**Fig.-4:**
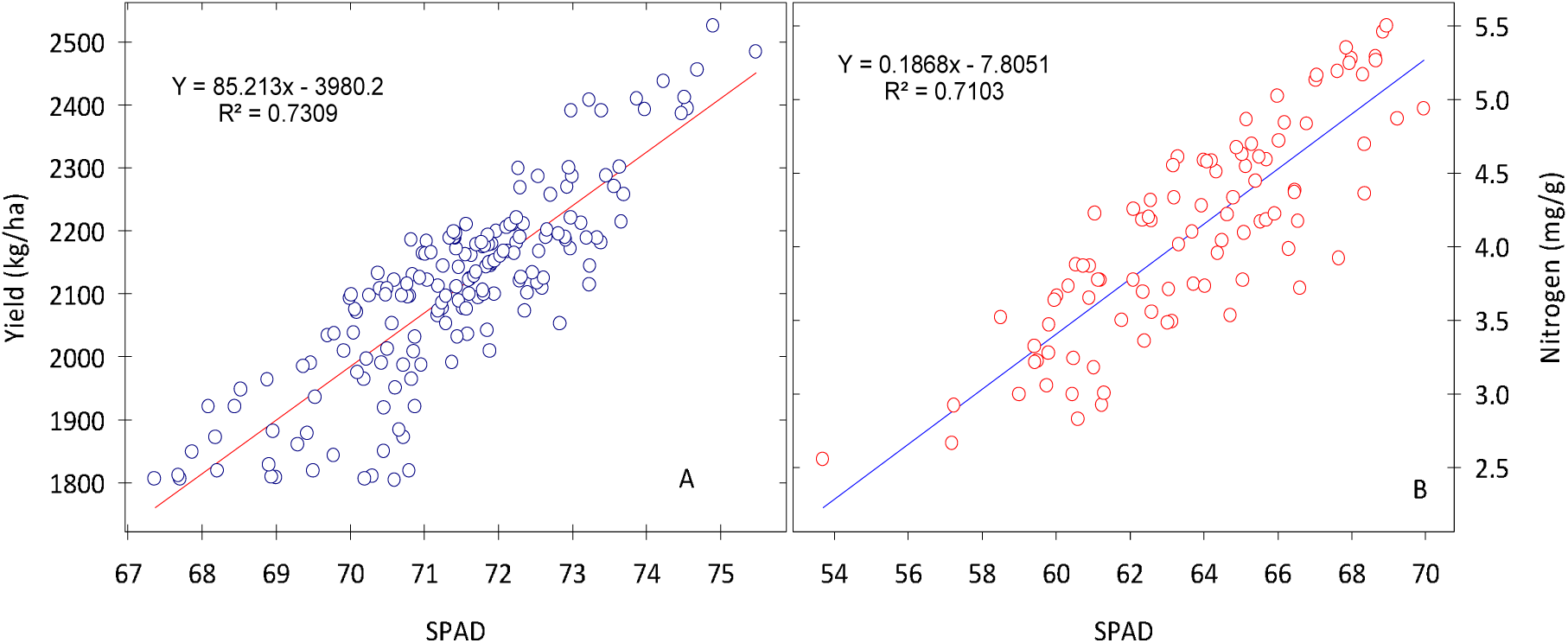
Relationship between SPAD values and yield (A); and SPAD with Nitrogen (B)

### Clonal Characterisation

The clonal signature of the UPASI released clones and an important estate selection was created to identifying the superior materials from the seedling blocks or infillings of an area of interest. The signature of the respective clones was established using a supervised image classification method and the algorithm was applied to the images of seedling fields. The plotrix of the UPASI released clones was depicted in the Fig.-5. From Fig.-6, it was observed that using supervised image segmentation algorithm can be exploited for the identification and composition of infilling clones in the area or some of the elite clones like materials from the seedlings block for crop improvement.

**Fig.-5:**
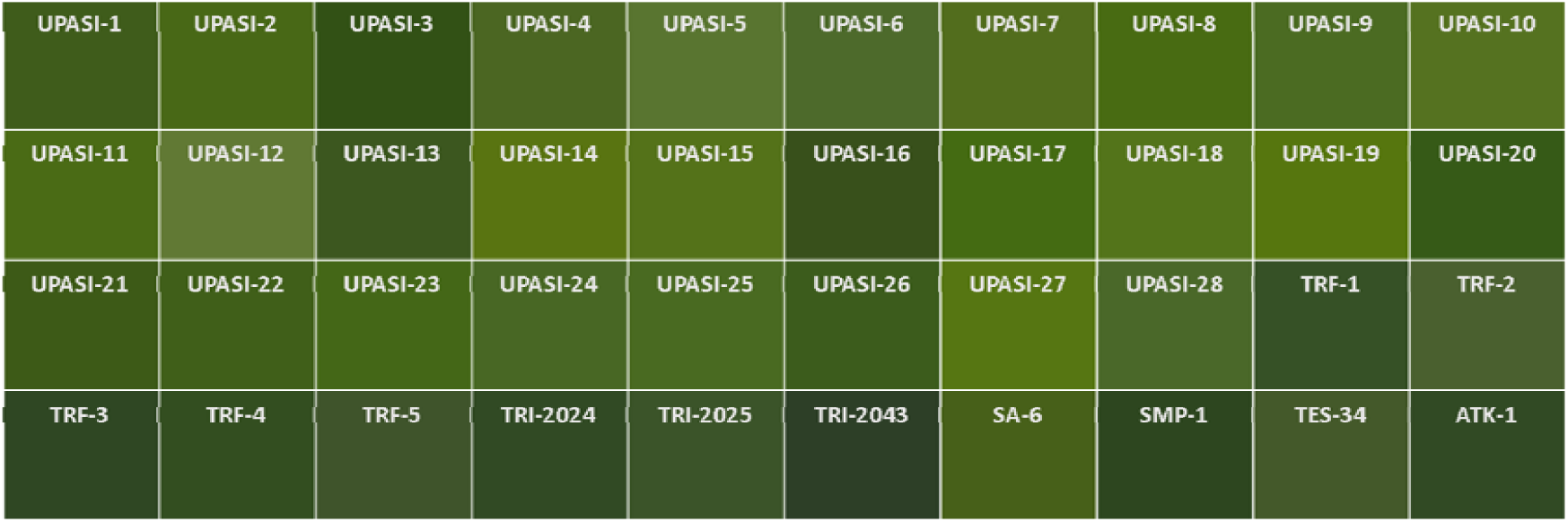
Colour profile of UPASI clones and estate selections.

**Fig.-6:**
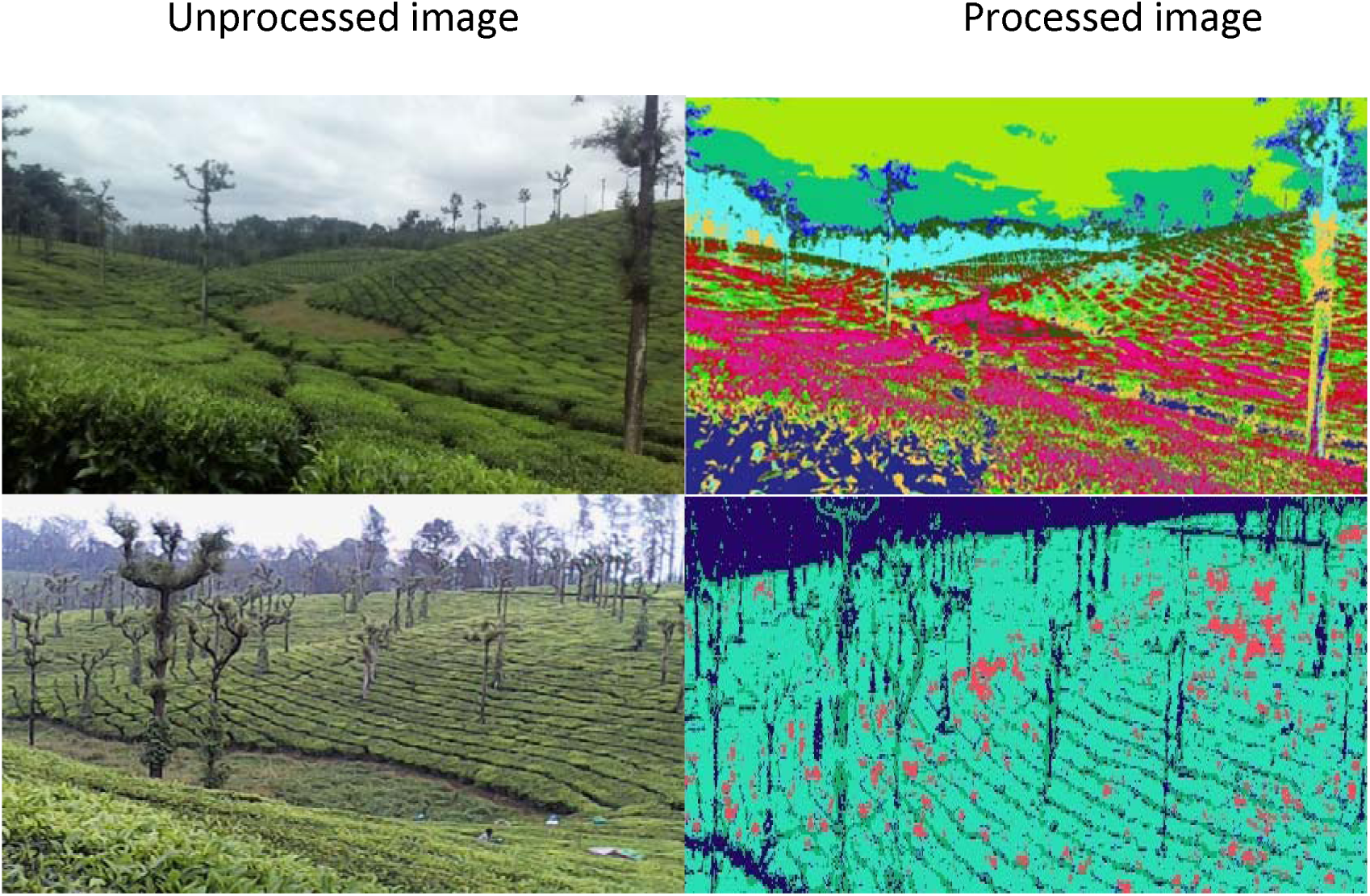
Application of supervised classification method for clonal identification (top row) and elite clones like materials in the seedling block (bottom row) using image segmentation analysis.

### Disease assessment by Image segmentation

For the assessment of blister blight disease, the images were subjected to K-mean hierarchical clustering and the area of infected portions was measured from the targeted cluster. Fig. 7 shows the original images followed by output segmented clusters revealed that the captured image can be classified into a different class of diseases. The classification of images with the Minimum Distance Criterion shows that there was about 95% accuracy in measuring the area of the target and the remaining 5% error consider as misclassification. Form Fig.-8, it was understood that inbuilt GPS of a smartphone can be utilised for plotting the disease occurrence with geographic information. This method can be used for monitoring disease dynamics before and after the application fungicides on a real-time basis.

**Fig.-7:**
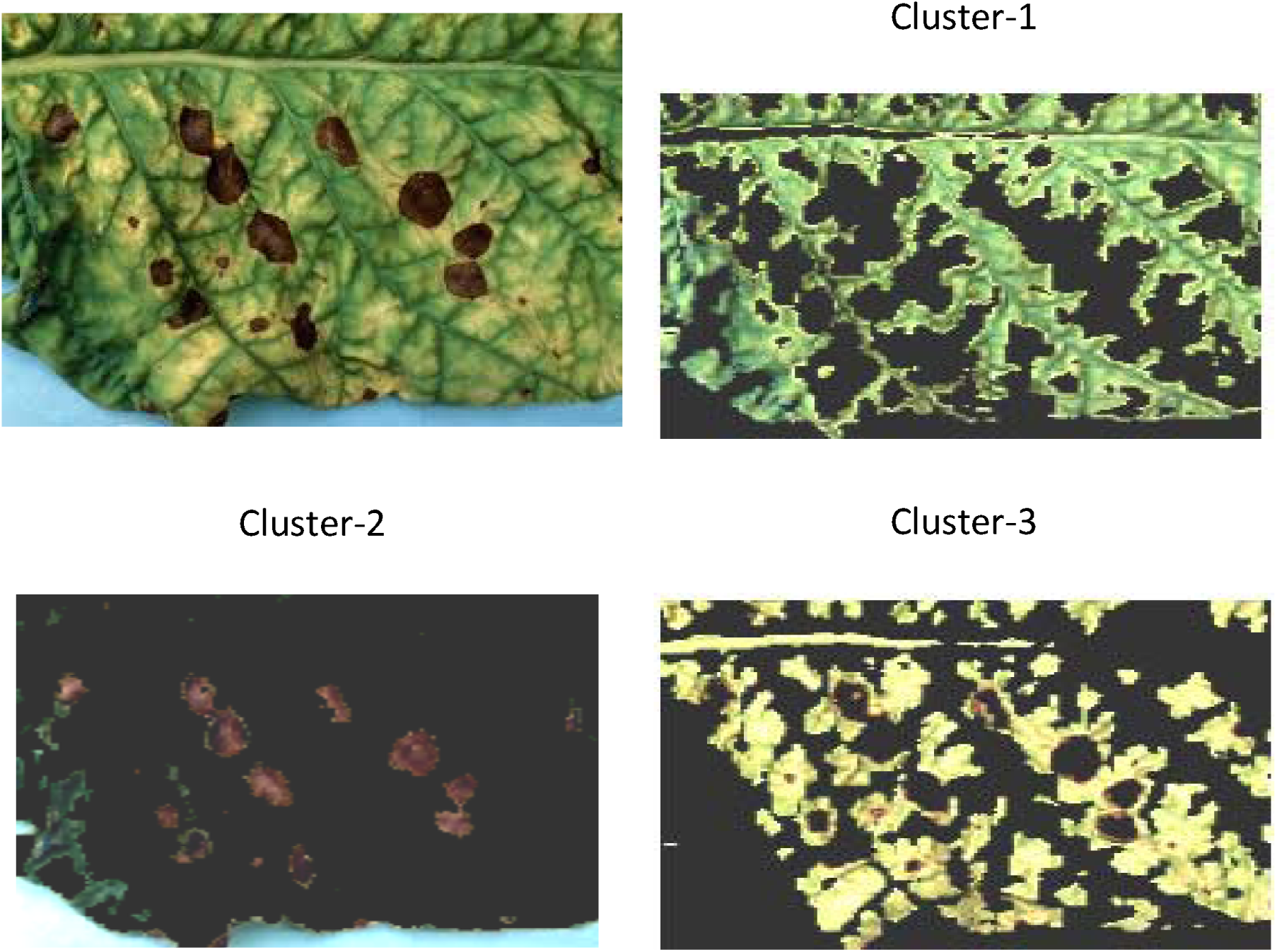
Application of image processing methods for disease assessment.

**Fig.-8:**
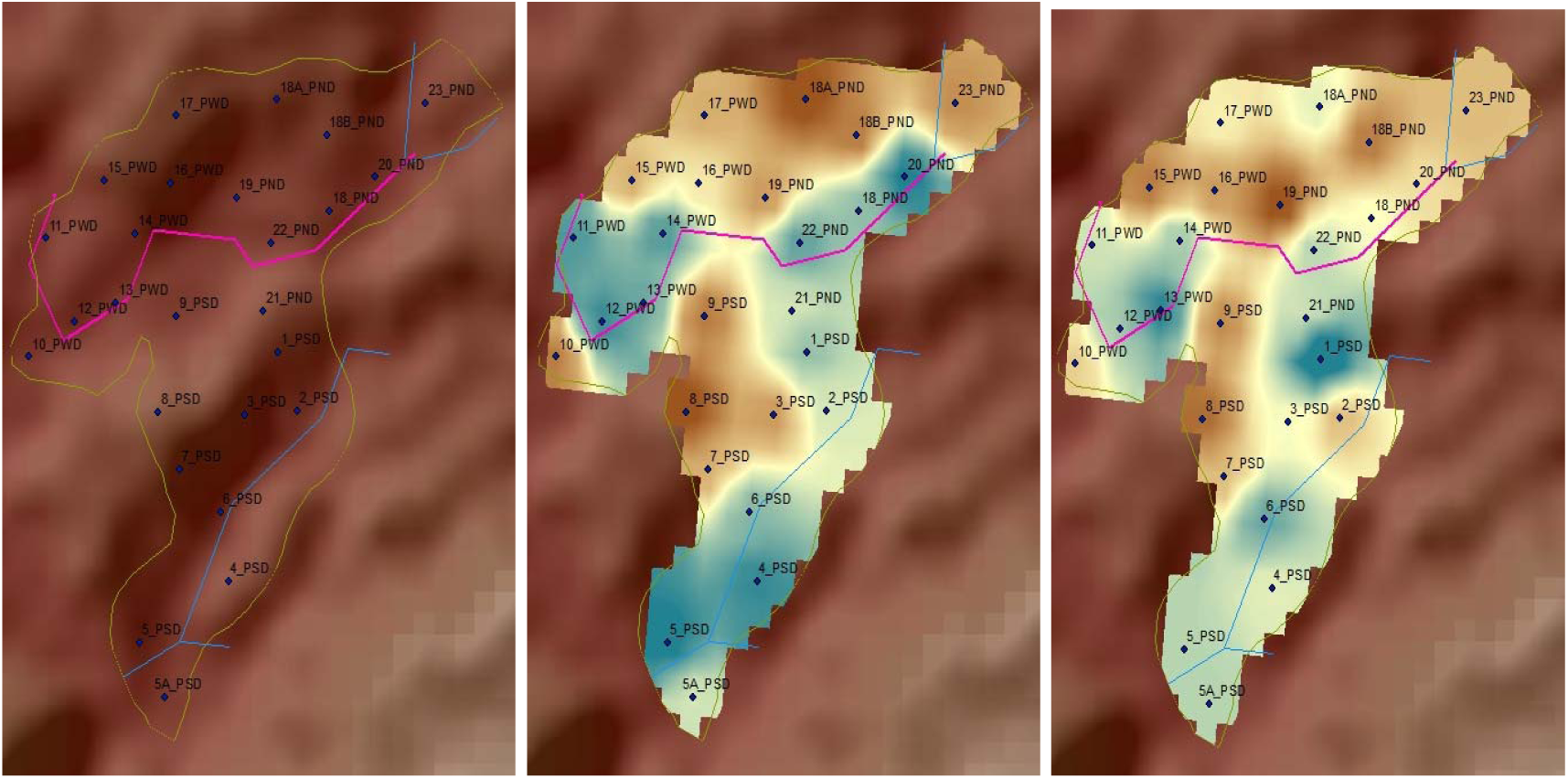
Spatial interpolation of blister disease (A) percentage before (B) and after chemical control (C).

## Conclusion

The study demonstrated various non-invasive image processing methods for the quantification of chlorophyll, nitrogen and disease. The methods can enable common agrarian for real-time estimation and assessment of nutrients and disease and its spatial dynamics for larger area management. Further study is required for validation and developing an Android application for planter’s utility.

